# Male genital lobe morphology affects the chance to copulate in *Drosophila pachea*

**DOI:** 10.1101/816538

**Authors:** Bénédicte M. Lefèvre, Diane Catté, Virginie Courtier-Orgogozo, Michael Lang

**Affiliations:** Team “Evolution and Genetics”, Institut Jacques Monod, CNRS, UMR7592, Université de Paris, 15 rue Hélène Brion, 75013 Paris

**Keywords:** *Drosophila pachea*, primary sexual trait, mating behaviour, mate-competition experiments, genitalia

## Abstract

**Introduction:** Male genitalia are thought to ensure transfer of sperm through direct physical contact with female during copulation. However, little attention has been given to their pre-copulatory role with respect to sexual selection and sexual conflict. Males of the fruitfly *Drosophila pachea* have a pair of asymmetric external genital lobes, which are primary sexual structures and stabilize the copulatory complex of female and male genitalia. We wondered if genital lobes in *D. pachea* may have a role before or at the onset of copulation, before genitalia contacts are made.

**Results:** We tested this hypothesis with a *D. pachea* stock where males have variable lobe lengths. In 92 mate competition trials with a single female and two males, females preferentially engaged into a first copulation with males that had a longer left lobe and that displayed increased courtship vigor. In 53 additional trials with both males having partially amputated left lobes of different lengths, we observed a weaker and non-significant effect of left lobe length on copulation success. Courtship durations significantly increased with female age and when two males courted the female simultaneously, compared to trials with only one courting male. In addition, lobe length did not affect sperm transfer once copulation was established.

**Conclusion:** Left lobe length affects the chance of a male to engage into copulation. The morphology of this primary sexual trait may affect reproductive success by mediating courtship signals or by facilitating the establishment of genital contacts at the onset of copulation.

## Introduction

Males and females exhibit different reproduction strategies due to higher energy costs of larger female gametes compared to smaller and more abundant male gametes [1]. This implies male-male intrasexual mate-competition for siring the limited female gametes. In turn, females may optimize reproduction by choosing males that confer survival and fecundity benefits to her and to the offspring. This dynamics has been formalized into genetic models [2, 3] involving female preferences for particular male characters and male mate competition, and can contribute to the rapid evolution of female preferences and male sexual attributes. There is also evidence for male mate-choice and female intra-sexual selection [4–6], showing that traditional sex-roles can change between species and in individuals under diverse conditions. In general, sexual selection implies a sexual conflict whenever male and female reproductive fitness strategies differ [7].

Across animals with internal fertilization, genitalia are usually the most rapidly evolving organs [8]. Several hypotheses propose that the evolution of genitalia is based on sexual selection and a consequence of sexual conflict between males and females at different levels of reproduction, including competition of sperm from different males inside the female (sperm competition) [9], female controlled storage and usage of sperm to fertilize eggs (cryptic female choice) [8, 10, 11], or sexually antagonistic co-evolution of aggressive and defensive morphologies and/or behaviors between male and female to control fertilization decisions [12, 13]. Male courtship is well known as a behaviour to attract females before copulation begins, but has also been reported to occur during or even after copulation [14, 15]. It is thought to be a widespread and key aspect of copulation in the cryptic female choice scenario, where males stimulate the female to utilize his own sperm [8, 10]. A variety of insect species were reported where males had evolved elaborated male genital structures that are apparently used for courtship throughout copulation to stimulate the female through tapping or other physical stimuli [14, 15].

Traditionally, sexual traits are categorized as primary or secondary. Genital structures are considered “primary” when they are directly used for the transfer of gametes during copulation or when they contribute to the complexing of female and male copulatory organs [8, 16]. Other traits that differ between sexes and that are linked to reproduction are considered “secondary” sexual traits. For primary sexual traits, sexual selection is traditionally thought to act during or after copulation [8, 15], while secondary sexual traits can be involved in pre-copulatory mate-choice via visual, auditory or chemical long range signalling [17–25].

Primary genitalia could also be possibly involved in pre-copulatory mate choice. Supposing that a particular male genital trait enhances or favours female fecundity, it might become preferred by the female during pre-copulatory courtship or at the moment of genitalia coupling [26–28]. This preference could rely on direct benefits to reduce energy investment or predation risk caused by unsuccessful copulation attempts, or on indirect male genetic quality to be inherited into the offspring generation. Several examples of primary genitalia used in pre-copulatory courtship signalling are known in vertebrates. For example, male and female genital display, swelling, or coloration changes are associated with sexual activity [29–31] and penile display was also observed in lizards [32]. Human male penis size was found to influence male attractiveness to women [33]. In diverse mammalian species, females have also evolved external and visible structures that resemble male genitals, a phenomenon called andromimicry [31, 34, 35]. These structures are likely used in visual signaling, so it is possible that the male analogous parts also do. Furthermore, females of some live-bearing fish species were reported to prefer to mate with males with large gonopodia [36, 37]. These cases involve visual stimuli and large genital organs. Only a few studies examined the roles of primary genitalia in invertebrates during courtship or at the onset of copulation before phallus intromission is achieved [26–28, 38, 39]. Longer male external genitalia of the water strider *Aquarius remigis* were associated with higher mating success [28]. In addition, male genital spine morphology of *Drosophila bipectinata* and *Drosophila ananassae* were reported to evolve in response to direct competition among males for securing mates in a so called ‘scramble competition’ scenario [26, 27]. In general, the role of primary sexual traits in establishing copulation or their role during courtship needs further investigation.

The fruitfly species *Drosophila pachea* is a promising model to study the evolution of primary sexual traits, especially with respect to the evolution of left-right asymmetry. Male *D. pachea* have an asymmetric aedeagus (intromission organ) and a pair of asymmetric external lobes with the left lobe being approximately 1.5 times longer than the right lobe [40–42] (Figure 1). The genital lobes have likely evolved during the past 3-6 Ma and are not found in closely related species [43]. *D. pachea* couples also mate in a right-sided copulation position where the male rests on top of the female abdomen with the antero-posterior midline shifted about 6-8 degrees to the right side of the female midline [41, 42, 44]. Male and female genitalia form an asymmetric complex during copulation and the asymmetric lobes stabilize this genital complex [44]. Furthermore, *D. pachea* is among the *Drosophila* species that produce the longest (giant) sperm [45, 46], and their ejaculates contain in average just about 40 sperm cells [47]. Thus, a particular right-sided mating position could potentially be associated with optimal transfer of giant sperm during copulation [42].

**Figure 1:**
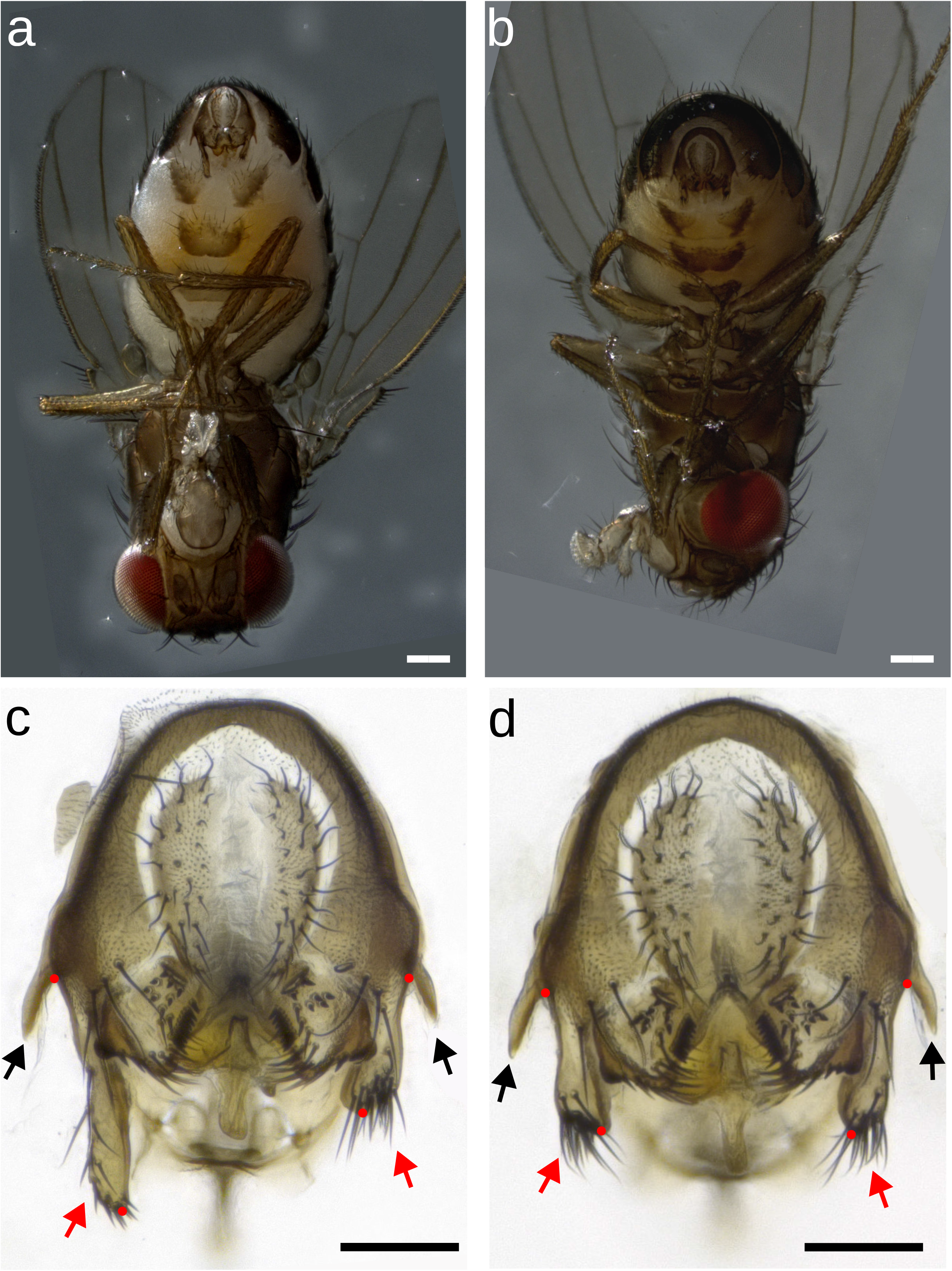
*Drosophila pachea* male genital lobes. (a) Male of stock 15090-1698.02 in ventral view and with asymmetric lobes. (b) Male of the selection stock with apparently symmetric lobes. The scale bars correspond to 200 μm. (c,d) Posterior view of a dissected male terminalia, (c) stock 15090-1698.02 with asymmetric lobes (d) selection stock. The scale bars correspond to 100 μm, dots indicate lobe length measurement points, red arrows point to the lobe tips and black arrows indicate the lateral spines.

The aim of this study was to test if the asymmetric genital lobes of *D. pachea* would have an effect on pre-copulatory mate-choice, in addition to their role in stabilizing the complex of male and female genitalia during copulation. Previously, we found that males originating from one of our *D. pachea* laboratory stocks possess short and rather symmetric lobes [41] (Figure 1), while others have the typical size asymmetry. The variability in left genital lobe length that we observed within this fly stock enabled us to test if lobe length might affect pre-copulatory courtship or mate competition. We selected *D. pachea* males with short left lobes and produced a stock with an increased variance of left lobe length. Next, we tested sibling males of this “selection stock” in mate-competition assays for their success to engage first into copulation with a single female. We also tested whether copulation success in our assay would be affected by male courtship vigour. Then, we surgically shortened the length of the left lobe in males that had developed long left lobes, to further test whether left lobe length affects copulation success. Finally, we assessed whether lobe length affects ejaculate allocation into female storage organs.

## Results

### Genital lobe lengths differ between *D. pachea* stocks

In our laboratory stock 15090-1698.01 (Additional datafile 1: Table S1), *D. pachea* males display a characteristic left-right size asymmetry of genital lobes with the left lobe being consistently larger than the right lobe (Figure 1a,c, Figure 2a, Additional datafile 1: Table S2, Additional datafile 2). Similarly, in stock 15090-1698.02, most males reveal larger left lobes, but a few individuals are observed with particularly small lobes that are rather symmetric in length (Figure 1b-d, Figure 2b). We selected for males with short lobe lengths and crossed them with sibling females for 50 generations (Methods). The resulting stock was named “selection stock”. We observed an increased variance of left lobe length compared to the source stock (Levene’s test: selection stock / 15090-1698.02: DF = 1/147, F = 11.506, P = 0.0008913) (Figure 2c, Additional datafile 2). In contrast, the right lobe length differed only marginally among the two stocks (Additional datafile 1: Table S2) and the variances of right lobe lengths were not significantly different among them (Levene’s test, DF = 1/147, F = 0.5164, P = 0.4735).

**Figure 2:**
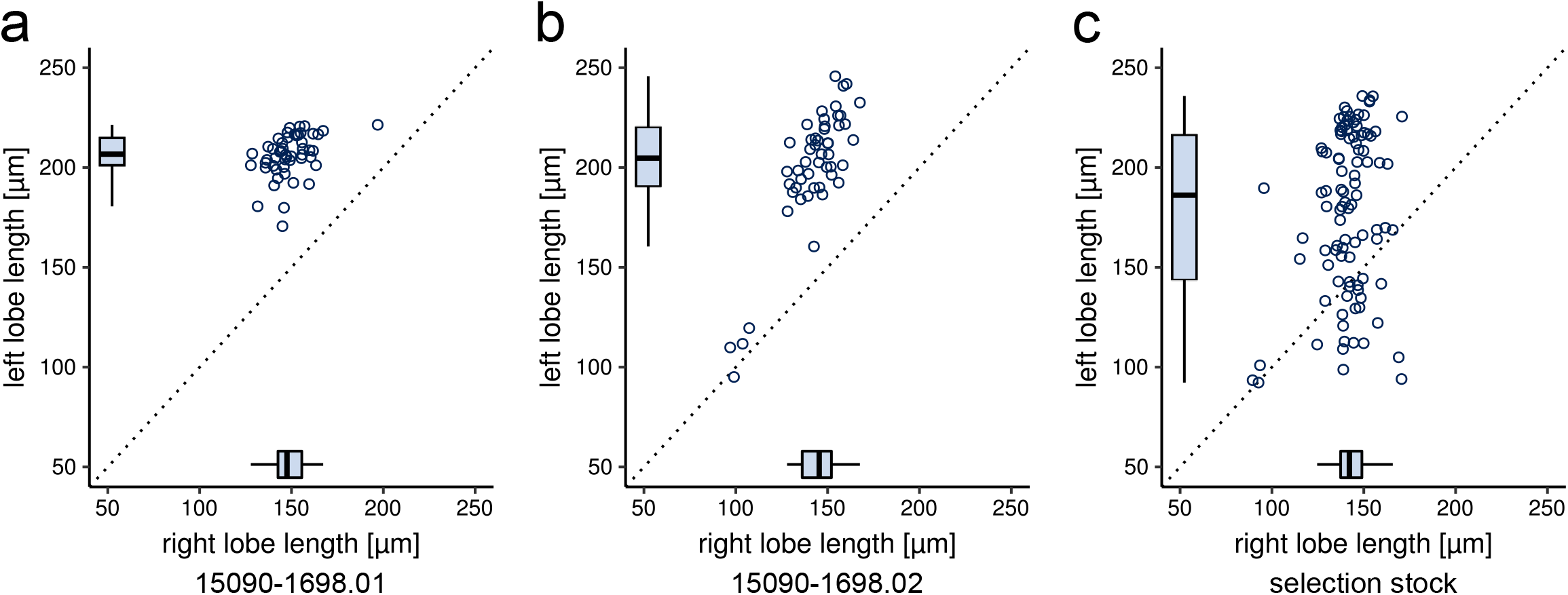
Genital lobe lengths differ between *D. pachea* stocks. Lengths of the left and right epandrial lobe lobes are presented for (a) stock 15090-1698.01, (b) stock 15090-1698.02, and (c) the selection stock. Sibling males of those used in the mating experiment are shown in panel (c). Each point represents one male. The variance of left lobe length is increased in the selection stock compared to the source stock 15090-1698.02 (Levene’s test: selection stock / 15090-1698.02: DF = 1/147, F = 11.506, P = 0.0009), while the variance of the right lobe length were not significantly different (Levene’s test, DF = 1/147, F = 0.5164, P = 0.4735).

### Male genital left lobe length, male courtship and failed copulation attempts have an effect on copulation success

We performed a mate competition assay (Methods) with two males and a single female to test if differences in genital lobe length and courtship behavior may influence the chance of a male to copulate first with a single female. This first engagement into copulation will be referred to as “copulation success”. We filmed courtship and copulation behavior (Additional datafile 1: Figure S1) of two males of the selection stock and a single female of stock 15090-1698.01 or 15090-1698.02. We annotated 111 trials where both males simultaneously courted the female (Methods) (51 and 60 trials with females of stocks 15090-1698.01 or 15090-1698.02, respectively). Then, we dissected the genitalia of both males to determine lobe lengths. We examined 54 additional trials with two courting males from the selection stock that had partially amputated left lobes and females of stock 15090-1698.01. This was done to test if other characters might co-vary with left lobe length and could potentially also have an effect on copulation success. We shortened the left lobes of both males in order to be able to neglect possible effects of the caused wound. The relative courtship activity or vigor of the two males in each trial was compared by quantifying periods that each male spent touching the female ovipositor or the ground next to it with the proboscis (mouth-parts), hereafter referred to as “licking behavior” (Methods). Licking was chosen over other behaviors because it occurs throughout courtship of *D. pachea* (Additional datafile 1: Figure S2) and was relatively easy to spot. We counted the sum duration of these periods (sum licking duration) and the number of licking periods (number of licking sequences). A copulation was considered to take place, once a male had mounted the female abdomen and achieved to settle into an invariant copulation position (Methods).

In the majority of trials (130/165), the first mounting attempt of a male resulted in a stable copulation. We did not observe any male behavior that could indicate an attempt of the non-copulating male to directly usurp the female as observed for *D. ananassae* or *D. bipectinata* [26, 27]. However, in 35 trials, a total of 51 failed mounting attempts were observed (Methods). These were not proportionally higher among trials with lobe amputated males (9 failed mounting attempts in 7 / 54 trials 13%) compared to unmodified males (42 failed mounting attempts in 28 / 111 trials 25%). Thus, the partial lobe amputation probably did not decrease the ability of a male to mount a female. Only 15/51 failed mounting attempts lasted 8 seconds or longer. Among those, the non-mounted male continued to court the female in 10/15 attempts. However, this was also observed in 131/165 successful mounting attempts and these proportions are not significantly different (χ^2^-test, χ^2^ : 1.4852, df = 1, P = 0.2230). Altogether, mounting failure was not found to be significantly associated with the courtship activity of the other male.

Different variables were tested to have en effect on male copulation success by using a Bradley Terry Model [48] (Methods). In trials with unmodified males, copulation success was positively associated with left lobe length and the sum licking duration of the males during courtship (Table 1). The effect of sum licking duration is illustrated when comparing the ratio of sum licking duration and total courtship duration between the copulating and the non-copulating male (Figure 3a). This ratio was higher in the copulating male compared to the non-copulating male. In particular, we identified a positive interaction of failed mounting attempts and female age, indicating that the negative effect of failed mounting attempts is pronounced in young females but not in older females. However, we observed failed mounting attempts in only 35 trials (Additional datafile 1: Figure S3) and the model estimates were sensitive to outlier observations (Additional datafile 1: Figure S10). Therefore, we relied on a simpler but more robust model that only takes sum licking duration and left lobe length into account (Table 1). More observations are required to evaluate the interaction of failed mounting attempts and female age.

**Table 1:** **Bradley–Terry (BT) model examining the effects of left lobe length, sum licking duration and failed mounting attempts on copulation success**, in 92 trials with unmodified males. Model: copulation success ~ sum licking duration + left lobe length, null deviance: 127.539 on 92 degrees of freedom, residual deviance: 98.416 on 90 degrees of freedom. Influential trials 87 and 98 were detected to have only marginal effects on model estimates.

| variable | estimate | std. error | z value | P (>\|z\|) |
| --- | --- | --- | --- | --- |
| sum licking duration | 0.714251 | 0.206293 | 3.462 | **0.000536** |
| left lobe | 0.007511 | 0.003254 | 2.308 | **0.020981** |

**Figure 3:**
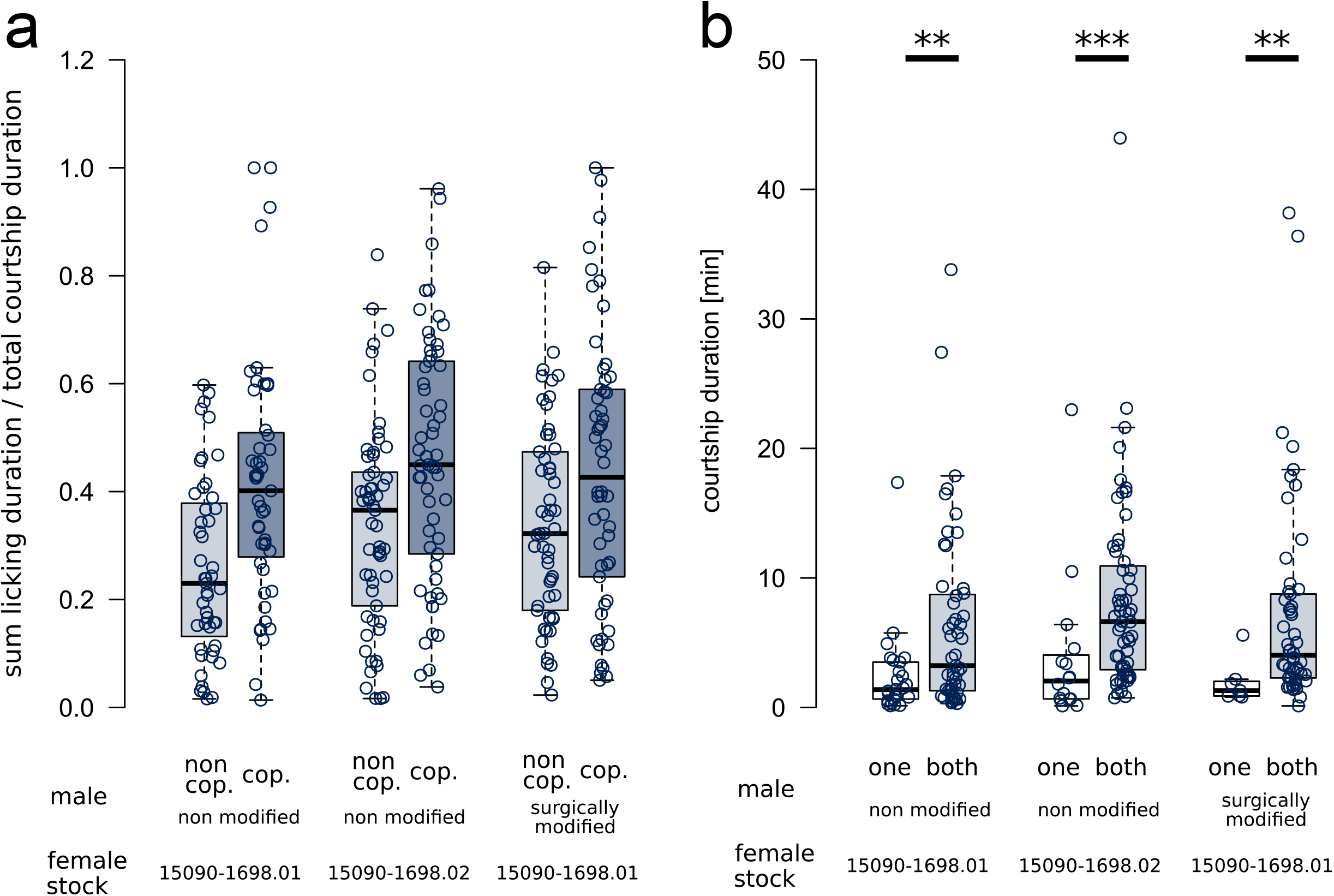
Courtship durations and relative courtship activity. (a) The ratio of sum licking duration and total courtship duration in each trial shows that the copulating male (cop, grey box-plots) had a higher average sum licking duration compared to the non-copulating male (non-cop, white box-plots). Each point represents one male. (b) The total courtship duration was shorter in trials with only one male displaying courtship signs (one, white box-plots) compared to trials with both males courting simultaneously (both, grey box-plots) (Mann-Whitney-Wilcoxon test, female stock 15090-1698.01, non modified males, W = 385, N = 25/51, P = 0.0053; females of stock 15090-1698.02, W = 219, N = 16/60, P = 0.0009; females stock 15090-1698.01, modified males, W = 62.5, N = 7/54, P = 0.0044). Horizontal bars indicate pairwise comparisons with P-values ** < 0.01 or *** < 0.001. Each point represents one mating experiment.

In trials with males that underwent lobe surgery, left lobe length was not significantly associated with copulation success (Table 2) or with male courtship behaviors. Thus, we cannot exclude the possibility that left lobe length co-varies with yet another unknown selected trait that reveals an advantage for the male to copulate. Since the copulating male still showed an increased sum licking duration compared to the non-copulating male (Figure 3), the non-significant model fit may thus reflect insufficient statistical power due to a smaller number of observations compared to trials with unmodified males.

**Table 2:** **Bradley–Terry (BT) model examining the effects of left lobe length, sum licking duration, and number of licking sequences on copulation success**, in 53 trials with males that had surgically modified left genital lobes. Model: copulation success ~ sum licking duration + left lobe length, null deviance: 73.474 on 53 degrees of freedom, residual deviance: 68.163 on 51 degrees of freedom

| variable | estimate | std. error | z value | P (>\|z\|) |
| --- | --- | --- | --- | --- |
| sum licking duration | 0.241981 | 0.166135 | 1.457 | 0.145 |
| left lobe | 0.007070 | 0.004615 | 1.532 | 0.126 |

### Courtship duration is increased when two males court a female simultaneously

Regardless of the stock for the female used, courtship durations appeared to be longer when both males courted the female simultaneously compared to trials where only one male courted (Mann-Whitney-Wilcoxon test, W = 1906.5, N = 48/165, P = 4.7×10^−8^) (Figure 3b). We also observed that courtship duration was strikingly increased in females that were older than 11 days (after emerging from the pupa) compared to younger females (Additional datafile 1: Figure S4) (Mann-Whitney test, W = 660, females younger than 12 days / females 12 days or older = 33 / 132, P *= 6*.333 × 10^−10^). Thus, older females may be less likely to copulate and may also discriminate less between males based on failed mounting attempts (see above).

### Lobe length is not associated with the amount of ejaculate in female sperm storage organs

To test whether lobe length influences the amount of ejaculate deposited into the female after copulation, we dissected a random subset of females (54 of stock 15090-1698.01 and 24 of stock 15090-1698.02) in trials with non-modified lobes that revealed a copulation. Neither right lobe length, left lobe length, sum licking duration or female age (days after emerging from the pupa) was significantly associated with ejaculate content in the spermathecae (Table 3). However, male adult age and copulation duration had a negative effect. Given that *D. pachea* males become fertile about 13 days after emerging from the pupa [49], our results indicate an optimal period of male ejaculate transfer at the beginning of their reproductive period. Copulation duration had a negative effect on sperm transfer (Table 3), indicating that failed sperm transfer may result in prolonged attempts and longer copulation duration.

**Table 3:** Male age and copulation duration affect spermathecae filling levels. Linear model (spermathacae filling level ~ sum licking duration + left lobe length + right lobe length + copulation duration + female age + male age), 32 observations were deleted due to missingness: sum licking duration 30 values, left lobe length 1 value), right lobe length 1 value, multiple R^2^: 0.4975, adjusted R^2^: 0.4201, F-statistic: 6.434 on 6 and 39 DF, P-value: 8.956e-05

| variable | F value | P |
| --- | --- | --- |
| sum licking duration | 0.8658 | 0.357838 |
| left lobe length | 0.0447 | 0.833711 |
| right lobe length | 2.4460 | 0.125901 |
| copulation duration | 11.3116 | **0.001737** |
| female age | 0.3300 | 0.568960 |
| male age | 23.6079 | **1.954e-05** |

## Discussion

### Left lobe length affects the chance of a male to copulate

We observed that most females copulated first with the male that had the longer left lobe when two males courted simultaneously in our trials. These results suggest that a long left lobe might not only stabilize an asymmetric genital complex during copulation [44], but also increases the chance of a male to engage into copulation. With our data, we cannot exclude that the sensing of the lobe length by the female occurs just before intromission, while male and female genitalia are contacting each other or if it is used even during courtship before genital contacts establish. Other factors, unrelated to lobe length, can also bias copulation. For example, females might prefer to copulate with males of larger body size or that display particular courtship behaviors. In our study, we found no significant effect of tibia length on copulation success, which was used as an approximation for overall body size. However, the mutation(s) associated with lobe length variation in this stock are unknown. It is possible that one or several alleles associated with lobe length still segregate in this stock, even after 50 generations of inbreeding. If such segregating mutations have pleiotropic effects on other male characters (morphological, physiological or behavioural), these traits are expected to co-vary together with lobe length and the effect of lobe length that we detected might actually be due to the other factors. In trials with artificially shortened left lobes , males were taken from the selection stock but had the expected wild-type lobe asymmetry. Left lobe length difference was then artificially introduced in those males, so that it would not be expected to co-vary with the relative phenotypic expression of the supposed underlying mutation. In such trials, left lobe length after artificial length reduction was not associated with copulation success so that we cannot rule out the possibility that left lobe length in our initial experiment with unmodified males affects copulation success indirectly via a pleiotropic mutation affecting both, lobe length and other traits. It is possible that we failed to detect a significant effect because of a reduced variation in left lobe length among the lobe-modified males, compared to the initial experiment with non-ablated males. In addition, we could not measure left lobe length on living males before lobe amputation. Therefore, we could not test whether the lobe length “before” length reduction influences male mating success. Furthermore, lobe amputation did not only cause a length reduction of the left lobe but also removed the bristles located at the distal tip. Laser ablation experiments have shown that these bristles contribute to the stabilization of the mating complex during copulation [44]. The bristles may also be important in pre-copulatory events, but we could not measure this since they were removed from both males in our trials. Future mate-choice experiments could be carried out with an unmodified male and a male where only those bristles are ablated [44].

### Male intra-sexual competition is potentially reflected by male courtship intensity

Previous analyses in *D. ananassae* and *D. bipectinata* relate male genital spines to pre-copulatory abilities to achieve a copulation due to increased male intrasexual competitiveness in so called “scramble competition” or vigorous copulation attempts with females that are optionally already copulating with anoher male [26, 27]. In *D. pachea*, this behavior was not observed and most mounting events of a male resulted in a stable copulation. Once a copulation started, the non-copulating male was never observed to separate the couple. Instead, it continued to display courtship behaviors such as licking, wing vibration or touching the female side in the majority of cases. In addition, male mounting was only observed upon female wing opening and oviscapt protrusion. This indicates that mounting in *D. pachea* is a complex behavior involving both sex partners. One possibility might be that upon mounting the chance of a *D. pachea* male to engage into copulation thus may largely depend on male-female communication rather than on male male competition.

Total courtship durations were increased in trials where both males courted the female simultaneously compared to trials with only one male courting. This indicates that females potentially require more time to accept one of the males for copulation in cases where two males court simultaneously. Alternatively, longer courtship durations might indeed rely on male-male competition, causing mutual disturbances in courtship display. We detected a strong association of copulation success with the sum licking duration, which we used as an approximation for overall courtship activity or vigor of each male. This is achieved by maintaining its position close to the female oviscape in competition with the other male, suggesting that licking behavior in part reports male-intrasexual conflict.

### Potential female benefits gained from choosing males with long left lobes

Female mate choice on male sexual traits was suggested to rely at least in part on the way a sexual trait is displayed or moved [50]. This implies that a certain quantity but also accuracy of locomotor activity would matter in courtship and might better reflect overall male quality and “truth in advertising” than a morphological character or ornament alone. Our data suggest that male settling upon mounting the female might be crucial for a successful copulation since we identified a negative effect of failed mounting attempts on copulation success and a positive interaction of failed mounting attempts with female age, possibly indicating that young females are more discriminative compared to older females with respect to male mounting performance. However, this effect was only based on 35 trials (Additional datafile 1, Figure S3) and model estimates were sensitive to outliers. Future work must further evaluate this trend. Female mate choice is hypothesized to be based on direct benefits, affecting fecundity and survival of the female, but also on indirect benefits that relate to genetic quality, fecundity and survival of the progeny [50, 51]. Left lobe length and male mounting performance might therefore influence female mate-choice through enhancing the quality of male courtship signals and by enhancing the coupling efficiency of male and female genitalia upon mounting. It remains to be resolved what these courtship signals could be and how they might be presented to the female. We did not observe direct contact of the female with male genitalia prior to the male mounting attempts at copulation start. A possible pre-copulatory signal mediated by genital lobes could therefore be visual or vibratory. During Drosophila courtship, males perform abdomen shakes (“quivers”) that generate substrate-borne vibratory signals [52]. Perhaps the quiver frequency or amplitude could be affected by the length of the left lobe, thus producing a modified vibration signal. Alternatively, asymmetric lobes might be associated with lateralized courtship. For example, the beetle species *Sitophilus oryzae and Tribolium confusum* were observed to perform left-biased copulation attempts, which led to higher mating success over right-biased males [53]. However, in our study the male always mounted the female from the rear, along the female midline. We also noted that the licking side upon mounting was most commonly the rear of the female abdomen, but changed frequently in response to female abdomen movements and due to the presence of the other male. Therefore, we did not identify any lateralized male pre-copulatory behavior. This is in contrast with the lateralized posture of *D. pachea* males later during copulation, with the male being shifted about 6-8° to the right side of the female abdomen [41, 42, 44].

Alternatively, left lobe length might potentially associate with direct copulation benefits for male and female by ensuring the efficiency of ejaculate transfer during copulation. Males of *D. pachea* produce giant sperm and previous estimates revealed that a *D. pachea* male transmits only about 44 ± 6 sperm cells per copulation [47], while the maximum female sperm storage capacity in the spermathecae was estimated to be much higher and to be about 304 sperm cells [47]. Wild-caught *D. pachea* females were found to contain sperm from at least 3-4 males based on spermathecae filling levels [47]. As asymmetric lobes stabilize the genital complex [44], they might have a positive effect on the amount of sperm transferred per copulation. However, we have not observed any effect of lobe length on ejaculate presence in the spermathecae after copulation. We note that we did not test for the presence of seminal fluid in the female outside of the spermathecae. Also, our quantification method was not precise and sample size was limited. Accurate direct sperm counts requires radio-labeling methods [47], which was not applicable in our experimental setup. In future approaches, improved sperm counts should be applied to test for a correlation between lobe length, female-male copulation complex stability and sperm quantity in female spermathecae.

Based on our data, we note that left lobe length affects the chance of a male to establish genital contacts that lead to a firm complex of female and male genitalia, congruent with previous observations in other insects [26–28, 54]. In particular, the male genital spines in *D. ananassae* and *D. bipectinata* were observed to play a role in pre-copulatory events but had no measurable influence later on with respect to sperm transfer or fecundity [26, 27, 54].

Interestingly, copulation duration negatively affected ejaculate presence in the female. This observation suggests that the couple separates once successful sperm transfer is achieved. However, not all copulations reveal sperm transfer. In occasions of failure the copulation duration may therefore be prolonged to further attempt to mate. Copulation duration varies largely among Drosophila species from fractions of a minute to up to about 2 h in *D. acanthoptera* [42, 49]. Copulation duration in *D. pachea* is thus comparatively long with respect to other species of the genus, pointing to presumably complex copulatory interactions affecting sperm transfer. Future studies must analyze the internal processes of sperm transfer during *D. pachea* copulation to better understand this process and its possible limitations.

Indirect paternal effects on the growth and survival of offspring could potentially play a role in *D. pachea* female choice for males with long left lobes. It has been demonstrated that mate choice based on particular male sexual characters correlates with offspring survival in diverse species [18, 20–23, 55]. However, our experiment did not allow us to test for these effects because females were sacrificed after the mating experiment and no progeny was obtained. We assessed ejaculate filling levels instead of the amount of female progeny because *D. pachea* females rarely lay eggs after a single insemination and about four copulations are necessary for a female to start oviposition [56]. To test for indirect effects, future studies should ideally use progeny males from wild caught *D. pachea* females and test if subtle lobe length variation in those males would correlate with offspring survival in single couple crosses. A similar approach was used by Hoikkala et al. (1998) [55], who found that *Drosophila montana* courtship song frequency of wild caught males correlated with the survival rate of the male’s progeny.

### Age of reproductive activities appear to be non-overlapping in sibling males and females

The amount of ejaculate present in the female spermathecae after a single copulation was negatively affected by the age of the copulating male. Similarly, it was previously found that the number of progeny in single couple crosses (presumably involving multiple copulations) was also dependent on male adult age in *D. pachea* [57]. Males in *D. pachea* need about 10-14 days at 25° C after emergence to become sexually mature [49, 56, 57]. This is related to adult testis growth and production of giant sperm [47]. It was shown that the relatively long time to reach male sexual maturity impacts the proportion of sexually active *D. pachea* adults (operational sex ratio), which is female-biased [49]. Our findings suggest that males potentially have a maximum fertility period at the beginning of their sexual maturity at approximately 13-16 days after emerging from the pupa. The detected reduction of transferred ejaculate in males older than 16 days implies an additional male reproductive limitation.

It was hypothesized that the delay in male sexual maturity might potentially lead to decreased sibling mating in *D. pachea* [56]. In *D. melanogaster*, sibling matings yield fewer progeny compared to crosses with unrelated individuals [57]. In our study, the female was not a sibling of the two males. Courtship durations were shortest in trials with “young” virgin females, between 6 and 11 days. Given that females reach sexual maturity at about 4 days after emerging from the pupa [49, 57], our observation suggests that young females engage more quickly into copulation compared to older females. Overall, female and male maximum reproductive activities appear to be non-overlapping among siblings of opposite sex and similar age. Such an effect can be relevant in the wild if a cactus rot is colonized by a single previously fertilized female.

## Conclusion

Longer male left genital lobes potentially enhance the probability of *D. pachea* males to engage into copulation, which is expected to confer a reproductive advantage over males with shorter left lobes. The chance of a male to copulate first with a female was also affected by the relative male courtship activity and potentially by mounting attempt failures, especially for females at the beginning of their reproductive activity. The evolution of left lobe length might not only relate to its previously known function as a stabilizing device during copulation, but also through pre-copulatory mate choice by increasing the chances to copulate and to increase the coupling efficiency of genitalia upon mounting. Lobes do not appear to impact sperm transfer in later copulatory events. In addition, reproductive ages are non-overlapping between the two sexes potentially decreasing the amount of sibling mating in this species.

## Methods

### Fly stock establishment and maintenance

Two *D. pachea* stocks 15090-1698.01 and 15090-1698.02 were retrieved from the San Diego Drosophila Species Stock Center (now The National Drosophila Species Stock Center, College of Agriculture and Life Science, Cornell University, USA) (Additional datafile 1: Table S1). Flies were maintained in 25 × 95 mm plastic vials containing 10 mL of standard Drosophila medium (60 g/L brewer’s yeast, 66.6 g/L cornmeal, 8.6 g/L agar, 5 g/L methyl-4-hydroxybenzoate and 2.5% ^v^/v ethanol) and a ~ 10 × 50 mm piece of bench protection sheet (Bench guard). As *D. pachea* requires 7-dehydrocholesterol for proper development [58–60], we mixed the medium of each vial with 40 μL of 5 mg/mL 7-dehydrocholesterol, dissolved in ethanol (standard *D. pachea* food). Flies were kept at 25°C inside incubators (Velp) with a self-made light installation for a 12 h light: 12h dark photo-periodic cycle combined with a 30-min linear illumination change between light (1080 lumen) and dark (0 lumen). We used males of stock 15090-1698.02 to generate a new stock with increased proportions of males with short gential lobes. For this, we chose 3 males with apparently symmetric (aberrant) genital lobes and crossed them with 3-4 sibling virgin females. We repeated the selection with the progeny for a total of 36 generations. Then, we discarded males with clearly visible asymmetric (wild-type) lobes from the progeny for another 14 generations to derive the final stock (selection stock).

### Virgin fly selection

Virgin flies at 0-1 d after emerging from the pupa were CO2 anaesthetised on a CO2-pad (INJECT+MATIC Sleeper) under a stereo-microscope Stemi 2000 (Zeiss), separated according to sex and maintained in groups of 20-30 individuals. Males and females were raised until reaching sexual maturity, about 14 days for males and 4 days for females at 25° C [49]. This allowed us to use virgin individuals in each mating experiment. Males were anaesthetised on the CO2 pad (see above), sorted according to lobe morphology (asymmetric and symmetric lobes) and isolated into single vials at least two days before each mating experiment took place.

### Mate-competition assay

We used a single virgin female of wild-type stock 15090-1698.01 or 15090-1698.02 and two sibling males of the selection stock that visually differed in lobe length when inspected with a binocular microscope (Additional datafile 1: Figure S5). This selection was done to increase the average pairwise difference of lobe lengths between the two males. Specimens were introduced into a white, cylindrical mating cell (Additional datafile 1: Figure S1) with a diameter of 20 mm, a depth of 4 mm and a transparent 1 mm Plexiglas top-cover. Optionally, mating cells were concave with a diameter of 20 mm and a depth of 4 mm at the center (Additional datafile 3: trials 1-16). Flies were transferred without CO_2_ anaesthesia using a fly aspirator: a 7 mm diameter silicone tube closed at the tip with cotton and a 1000 μL wide bore (3 mm) micro-tip. Movie recordings were started as soon as the chamber was immediately put under the camera (see below).

All mating trials were carried out inside a temperature and humidity controlled climate chamber at 25° C ± 0.1° C and 80% ± 5% or 60% ± 5% (trials 1-15) humidity. We used digital cameras MIRAZOOM MZ902 (OWL) or DigiMicro Profi (DNT) to record copulation and courtship behaviour. The MIRAZOOM MZ902 (OWL) camera was mounted on a modified microscope stand (191348, Conrad) equipped with a 8-cm LED white light illumination ring (EB-AE-COB-Cover, YM E-Bright) and a platform to hold four individual cylindrical mating cells (Additional datafile 1: Figure S1). Trial 16 (Additional datafile 3) was filmed with the OWL camera and a concave shaped mating cell that was put on a flat plastic cover on top of the microscope stand. For trials 1-15, we used concave shaped mating cells that were put onto the stand of the DigiMicro Profi (DNT) camera. Data was acquired with the programs Cheese (version 3.18.1) (https://wiki.gnome.org/Apps/Cheese) or GUVCVIEW (version 0.9.9) GTK UVC (trials 1-15) on an ubuntu linux operating system in webm or mkv format. Up to four trials were recorded in a single movie, which was split after recording and converted into mp4 format with Avidemux 2.6 (http://www.avidemux.org/) to obtain a single movie per experiment.

We waited for a copulation to take place between one of the males and the female (see below). The mating cell was then shortly recovered from the climate chamber in order to remove the non-copulating male with an aspirator and to transfer it into a 2-mL reaction tube filled with 70% ethanol. This was performed after the copulating male appeared to have settled into a stable mating posture on the female abdomen, which was controlled by live-imaging. The copulating male and the female were isolated from the mating cell after copulation had ended and were also isolated into single 2-mL reaction tubes filled with 70% ethanol. Optionally, the female was kept alive for 12-24 h in a vial containing 5-10 mL grape juice agar (24 gr/L agar, 26 gr/L sucrose, 120 mg/L Tegosept, 20% ^v^/v grape juice, and 1.5% ^v^/v ethanol). The presence of eggs on the plate was systematically checked but never observed. Females were finally sacrificed to prepare spermathecae.

### Left lobe surgery

Epandrial lobe surgery was done on 5-6 day-old *D. pachea* adult males of the selection stock following Rhebergen et al. (2016). Males were anaesthetised on a CO2 pad (see above) and then further immobilized with a small copper wire, which was slightly pressed onto the male abdomen. The left epandrial lobe was shortened to various lengths with micro dissection scissors (SuperFine Vannas Scissors, World Precision Instruments). Flies were let to recover for at least 7 days on standard *D. pachea* food at 25° C in groups. No mortality was observed in males that underwent lobe surgery, similar to what was observed by Rhebergen et al. (2016). Partially amputated lobes ranged in length from lacking at least their most distal tip to more reduced stumps whose length approximated the corresponding right lobe.

### Adult dissection and imaging

Adults were dissected in water with forceps (Forceps Dumont #5, Fine Science Tool) inside a transparent 25 mm round dissection dish under a stereo-microscope (Zeiss Stemi 2000). Male genitalia were isolated by piercing the abdomen with the forceps between the genital arch and the A6 abdominal segment and thereby separating the genitalia from the abdomen. We also dissected the left anterior leg of each dissected male. Female spermathecae were recovered after opening the ventral abdomen with forceps and removal of the gut and ovaries. The spermathecae were isolated and separated according to left and right sides and immediately examined using a microscope (see below). All dissected tissues were stored in 200 μL storage solution (glycerol:acetate:ethanol, 1:1:3) at 4° C.

For imaging, male genitalia were transferred into a dissection dish filled with pure glycerol and examined with a VHX2000 (Keyence) microscope equipped with a 100-1000x VH-Z100W (Keyence) zoom objective at 300-400 fold magnification. Genitalia were oriented to be visible in posterior view. The left and right lateral spines, as well as the dorsal edge of the genital arch were aligned to be visible in the same focal plane. In some preparations, the genital arch broke and lobes were aligned without adjusting the position of the dorsal genital arch. The experiment was discarded from analysis in cases where lobes could not be aligned. Male lobe lengths were measured on acquired images as the distance between the base of each lateral spine and the tip of each lobe (Figure 1c,d) using ImageJ version 1.50d (https://imagej.nih.gov/ij). Male legs were put on a flat glycerol surface with the inner side of the tibia facing to the camera. Legs were imaged at 200 fold magnification with the VHX 2000 microscope (Keyence) as described above.

We prepared the female paired sperm storage organs (spermathecae) at 12 h - 24 h after copulation (54 females from stock 15090-1698.01 and 24 females from stock 15090-1698.02, additional file 2) and examined the presence of ejaculate. *D. pachea* has giant sperm [47], which makes direct sperm counts difficult. Therefore, we determined an apparent average spermathecae filling level per female based on visual inspection, similar to Jefferson (1977) [57]. Female spermathecae were arranged on the bottom of the transparent plastic dissection dish, filled with water. Images were acquired using transmission light in lateral view with the Keyence VHX2000 microscope at 400-500 fold magnification. Ejaculate was directly visible inside the transparent spermathecae. Ejaculate filling levels were annotated for each spermatheca separately to match three different categories: 0, 1/6 or 1/3 of its total volume. Then, the average filling level for both spermathecae was calculated for each female.

### Annotation of courtship and copulation behaviour

Videos were analysed with the OpenShot Video Editor software, version 1.4.3 (https://www.openshot.org) to annotate the relative timing of courtship and copulation in our experiments (Additional datafile 1: Figure S2, Additional datafile 3). Data was manually entered into spreadsheets. We annotated the beginning of male courtship as the start of at least three consecutive male courtship behaviours according to Spieth (1952) [61], such as male touching the female abdomen with the forelegs, wing vibration, male following the female, and male licking behavior (see above). We estimated licking behavior throughout courtship of each male in trials were both males courted the female simultaneously. Licking occurs abundantly throughout courtship of *D. pachea* (Additional datafile 1, Figure S3) and was therefore chosen to assess the relative courtship contribution of the two males. We calculated the sum duration of all licking events per male and trial and the number of licking sequences. However, this behavior could not be quantified in 7 trials because the males changed positions very fast so that they could not be unambiguously distinguished (Additional datafiles 3,4). In all analysed trials, licking behaviour was observed throughout *D. pachea* courtship from shortly after courtship start to the start of copulation (Additional datafile 1: Figure S2). Both males reached at least once the licking position in all trials (Additional datafile 1: Table S3). This indicates that the female had physical contact with the proboscis of both males before copulation started (Additional datafile 1: Figure S2).

The beginning of copulation was defined as the moment when the male mounted the female abdomen. However, we only scored a copulation start if the male also settled into an invariant copulation posture. Upon mounting, the male moves its abdomen tip forth and back along the female oviscape. This behavior lasts for up to about 3 minutes and during this time the male grasps the female wings and abdomen with its legs. This behavior was previously described as settling period and ends with a characteristic right-sided, invariant copulation posture [41, 42, 44]. Any copulation that ended within the first minutes after mounting without settling was considered to be a “failed mounting attempt” (Additional datafile 4). The end of copulation was considered as the moment when male and female genitalia were separated and the male had completely descended with the forelegs from the female abdomen.

Copulation duration (Additional datafile 1: Table S3) was comparable to previous analyses of *D. pachea* copulation durations [41, 44, 47, 57]. We observed that copulation ended upon isolation of the non-copulating male in 3 trials with females of stock 15090-1698.01 and in 4 trials with females of stock 15090-1698.02, probably indicating premature copulation end due to specimen handling. However, the estimates were included into the dataset and had little impact on average copulation duration when comparing mean and median values.

### Mate-competition analysis

Our aim was to compare at least 50 mate-competition trials, where both males courted the female simultaneously and where copulation was observed. In total, we carried out 98 and 89 trials with females of stocks 15090-1698.01 and 15090-1698.02, respectively (Additional datafile 1: Table S2). We performed another 62 trials with males that underwent surgical manipulation of the left lobe and females of stock 15090-1698.01 (Additional datafile 1: Table S4). In total, we removed 36 trials from the analysis because either copulation was not observed until 1 h after recording started (21 trials), flies escaped, got injured or died inside the mating cell (4 trials) or the dissections of male genitalia failed (11 trials, Additional datafile 1: Table S4). We thus examined 152 trials with unmodified males and 61 trials with males that had undergone genital surgery, (see below, Additional file 1: Table S4). Among those, we observed that both males courted the female simultaneously in 111 and 54 trials, respectively. Statistical assessment with the Bradley Terry Model (see below) further required a complete dataset so that we excluded 20 additional trials with missing values, 19 trials with unmodified males (left lobe length: 2 trials, right lobe length: 2 trials, licking behavior and/or failed mounting attempts: 7 trials, tibia length: 8 trials) and one trial with lobe-modified males (no information on tibia length). In total, we considered 92 trials with unmodified males and 53 trials with lobe-modified males.

Data analysis was performed in R version 3.6 [62]. Male copulation success (see above) was evaluated with a Bradley Terry Model [48] using the R package BradleyTerry2 [63]. This model is suitable for logit fits to pairwise comparison data since it takes into account the special non-random (pairwise) structure of the dataset. We incorporated data (Additional dtafile 3) into a list with three objects (Pacdata, available on DRYAD): 1) “contest” with the pairings of males, a trial ID and a factor indicating trials with males that had modified left genital lobes, 2) trial-specific variables “conditions” containing female age, female stock, courtship duration and a factor indicating which male initiated courtship, 3) male-specific predictor variables “male_predictors” with left and right genital lobe lengths, male tibia length, sum licking duration, number of licking sequences and failed mounting attempts. We developed the statistical model (Additional datafile 1: Tables S5-11) by evaluating the effect of different variables on copulation success and by comparing pairs of nested models, using Chi-square tests with the generic anova() function [63]. Outlier observations were considered when the standardized model residuals were greater than 2. The effect of corresponding trials on model fits was evaluated by comparison to additional model estimates without these outlier trials. We estimated correlations of male specific predictor variables and among numeric trial specific variables with the corrplot R package (Additional datafile 1: Figure S6). We found a significant correlation of the male predictor variables ‘sum licking duration’ and ‘number of licking sequences’. In addition, the trial specific variables ‘female age’ and ‘courtship duration’ were significantly correlated (Additional datafile 1: Figure S4). Correlated parameters were not incorporated into the same model for assessing the best fitting model (Additional datafile 1: Tables S5-S12). We tested trial specific variables in interaction with all male-specific variables, but only found an interaction of female age with failed mounting attempts in the dataset with unmodified males (Additional datafile 1: Figure S4, Table S8, Table 1). We also tested for interactions between male specific predictor variables but did not find any. The best fitting model for the datasets was: copulation success ~ sum licking duration + left_lobe length.

Spermathecae filling levels were evaluated with a linear model using the lm() function: spermatheca filling level ~ left lobe length + right lobe length + sum licking duration + Number of licking sequences + copulation duration + female age + male age, and significance of each variable was evaluated with an F-test using anova().

## Supporting information

Additional datafile 1

Additional datafile 2

Additional datafile 3

Additional datafile 4

## List of abbreviations

Ma: million years ago
DF: degree of freedom
N: number of observations
P: probability

## Declarations

### Ethics approval and consent to participate

“Not applicable” --- This research focuses on invertebrate insects. There are no ethical considerations mentioned for these species according to EU Directive 86/609-STE123.

### Consent for publication

“Not applicable”

### Availability of data and material

The data sets supporting the results of this article are available in the DRYAD repository, https://datadryad.org/stash/share/jzbHRTO70QV55ejT_BxIcopEoG-wVDjRI_HpQK0ergY

### Authors’ contributions

ML and BL designed the experiments. BL and DC recorded fly copulation and performed light microscopy analysis of *Drosophila* male genitalia. BL and ML analysed the data. ML, VCO and BL wrote the manuscript. All authors have read and approved the manuscript.

## Acknowledgements

We thank Therese A. Markow for discussions. We are very grateful to Alexandre Peluffo for help with the statistical analysis, Claus Lang for technical support and Juliette Royer for dissections of *D. pachea* male genitalia.

## Supporting Information

**Additional datafile 1:** Supplementary Figures and Tables, **Figure S1:** Camera device for movie recording, **Figure S2:** Courtship and copulation duration in competition mating experiments, **Figure S3:** Failed copulation attempts compared to female age, **Figure S4:** Courtship duration increases with female age, **Figure S5:** Pairwise lobe length differences of males used in the competition mating experiment, **Figure S6:** Pearson correlations of male specific predictor variables, **Table S1:***Drosophila pachea* resources, **Table S2:** Genital lobe lengths in *D. pachea* stocks, **Table S3:** Courtship duration, male licking behavior and copulation duration in trials where both males courted the female, **Table S4:** Competition mating trials used for courtship analysis, **Tables S5-S12:** Bradley Terry Model Fit.

**Additional datafile 2:** Raw measurements of lobe lengths in different *D. pachea* stocks

**Additional datafile 3:** Measurements of mating behaviour, genital lobe lengths and spermathecae filling levels of individuals used in the competition mating experiments.

**Additional datafile 4:** Dataset for male licking behavior and male mounting.

