## Additional datafile 1 for "Male genital lobe morphology affects the chance to copulate in *Drosophila pachea*"

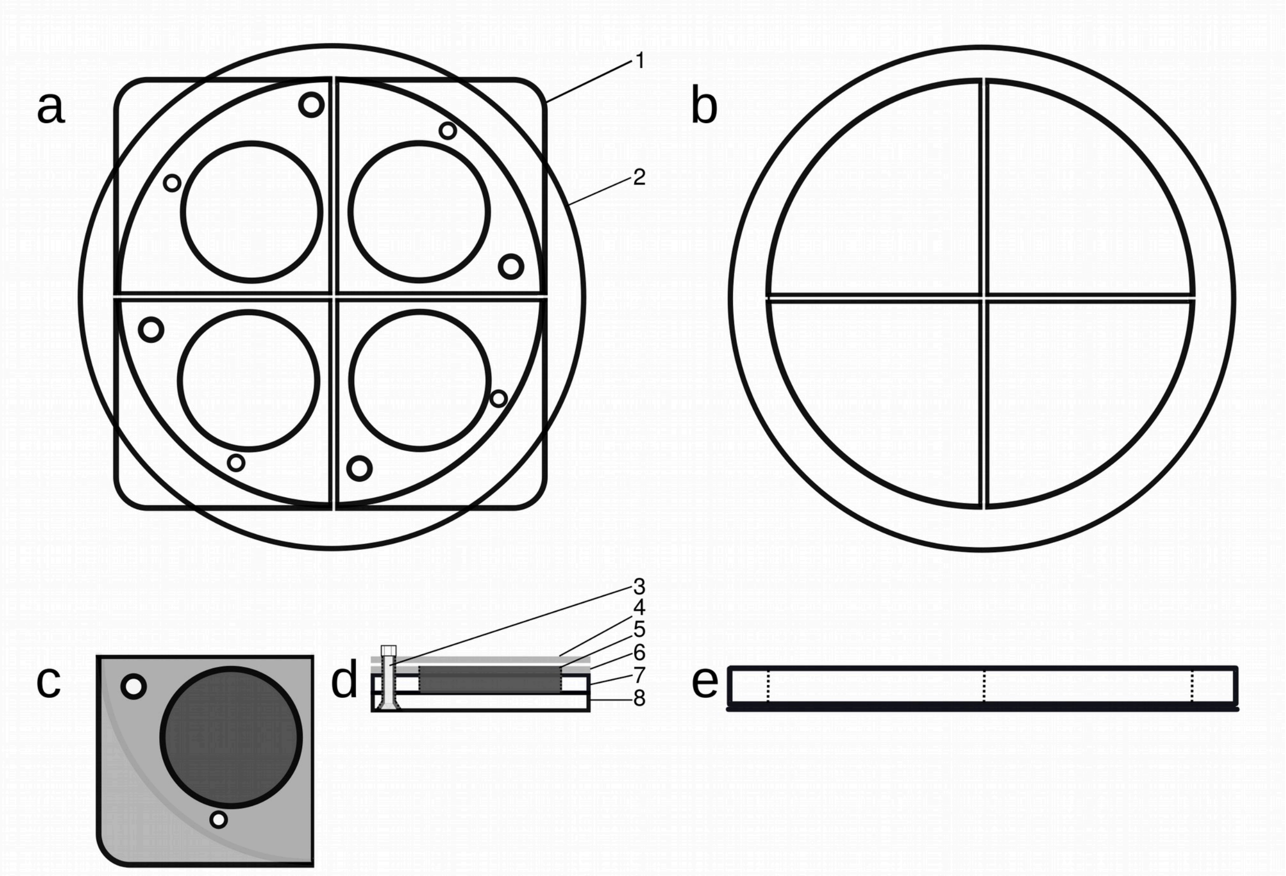


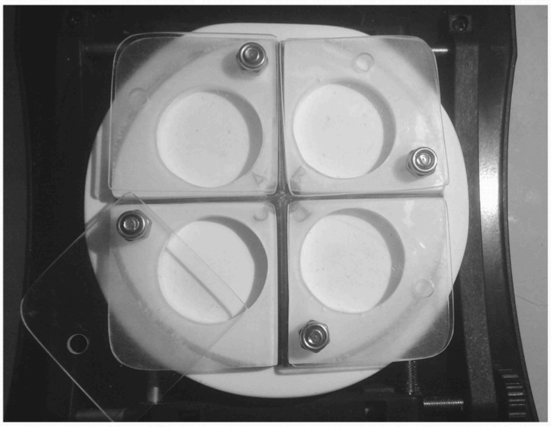


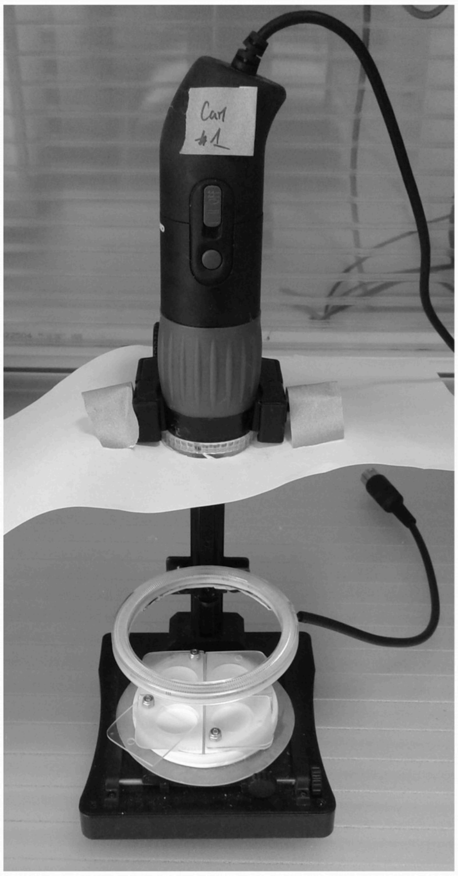


**Figure S1: Camera device for movie recording.** a) Schematic drawing of 4 mating cells (1) arranged on a round support (2); b) The round support has an inner diameter of 62 mm and is cross-divided (1 mm plexiglas) to hold four individual mating cells; c) the mating cell is a 30 mm quadrant (top view). A 3 mm screw (3) holds a transparent top cover (4), which has 3 mm bore for the screw and a 2 mm bore to introduce flies. d) The mating cell in lateral view shows the layer construction: It is assembled with a 3 mm screw (3) that holds the 1 mm thick transparent Plexiglas cover (4) on top of a 7 mm thick plastic unit with the cylindrical 20 mm x 4 mm inside of the mating cell (5). It consists of a 1 mm transparent Plexiglass layer (6) that serves as a handle and has an inner bore of 20 mm, a 3 mm thick white Plexiglas layer (7) with the inner bore, and a 3 mm thick white Plexiglas bottom layer (8). e) The round mating cell support in lateral view. It consists of a Plexiglas ring of 5 mm, an outer diameter of 70 mm and an inner diameter of 60 mm. It is glued to a 1 mm thick Plexiglas bottom layer. Dashed lines indicate the inner edge of the Plexiglas ring and the 1 mm cross division. f) The assembled mating cell device with 4 mating cells and the support mounted on a microscope stand (191348, Conrad). g) The mating cell device mounted on the microscope stand and equipped with a LED white light 3-12 V, 8-cm illumination ring (EB-AE-COB-Cover, YM E-Bright) and a camera (MIRAZOOM MZ902, OWL).

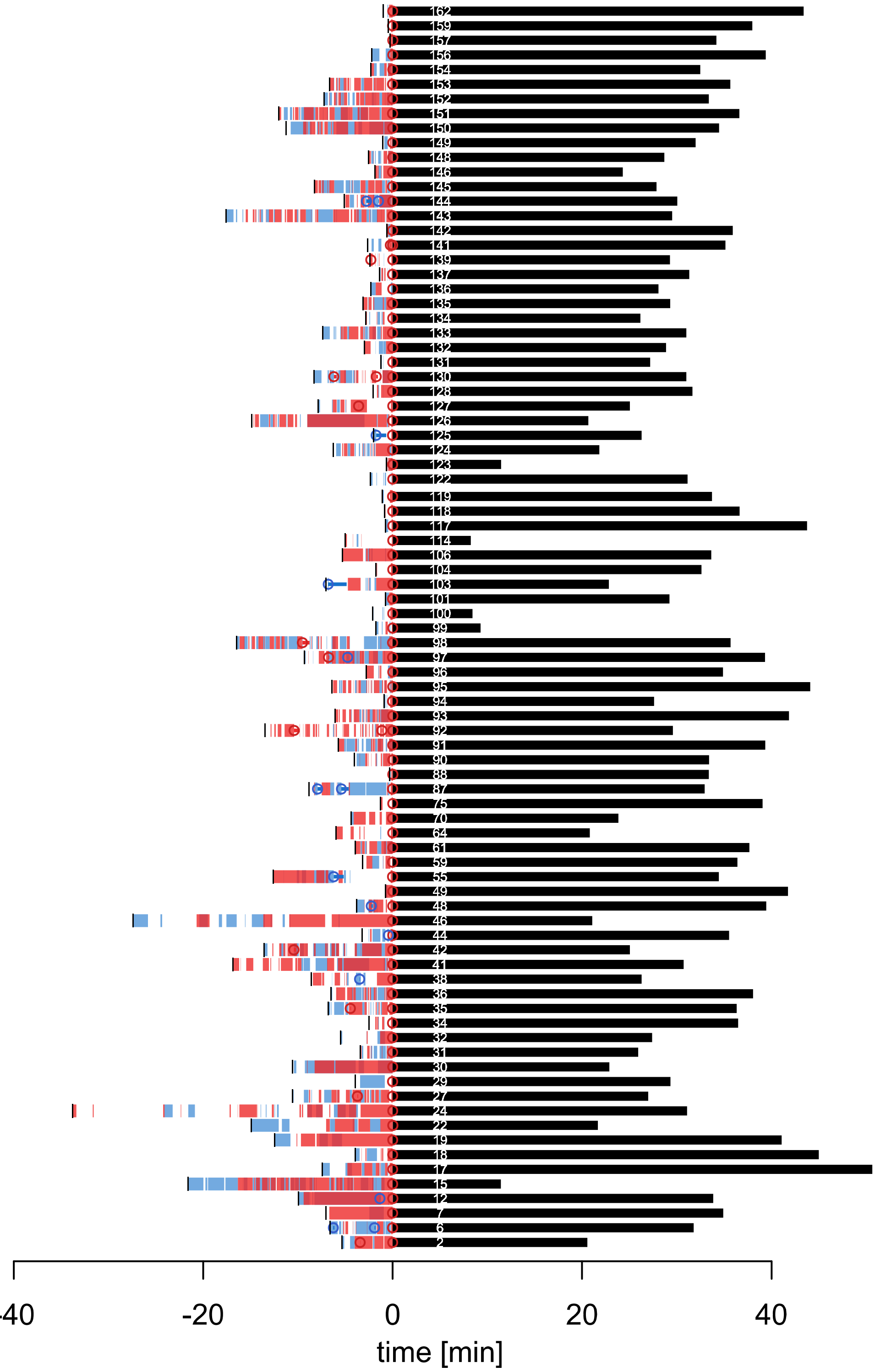


**Figure S2: Courtship and copulation duration in competition mating experiments.** The black dash indicates courtship start, segments indicate licking periods, open circles mark mounting events and horizontal bars indicate the durations of failed mounting attempts of the copulating male (red) and the non-copulating male (blue). The black bar indicates copulation duration and the number refers to the trial (mating ID), described in Additional datafile 3 and Additional datafile 4.

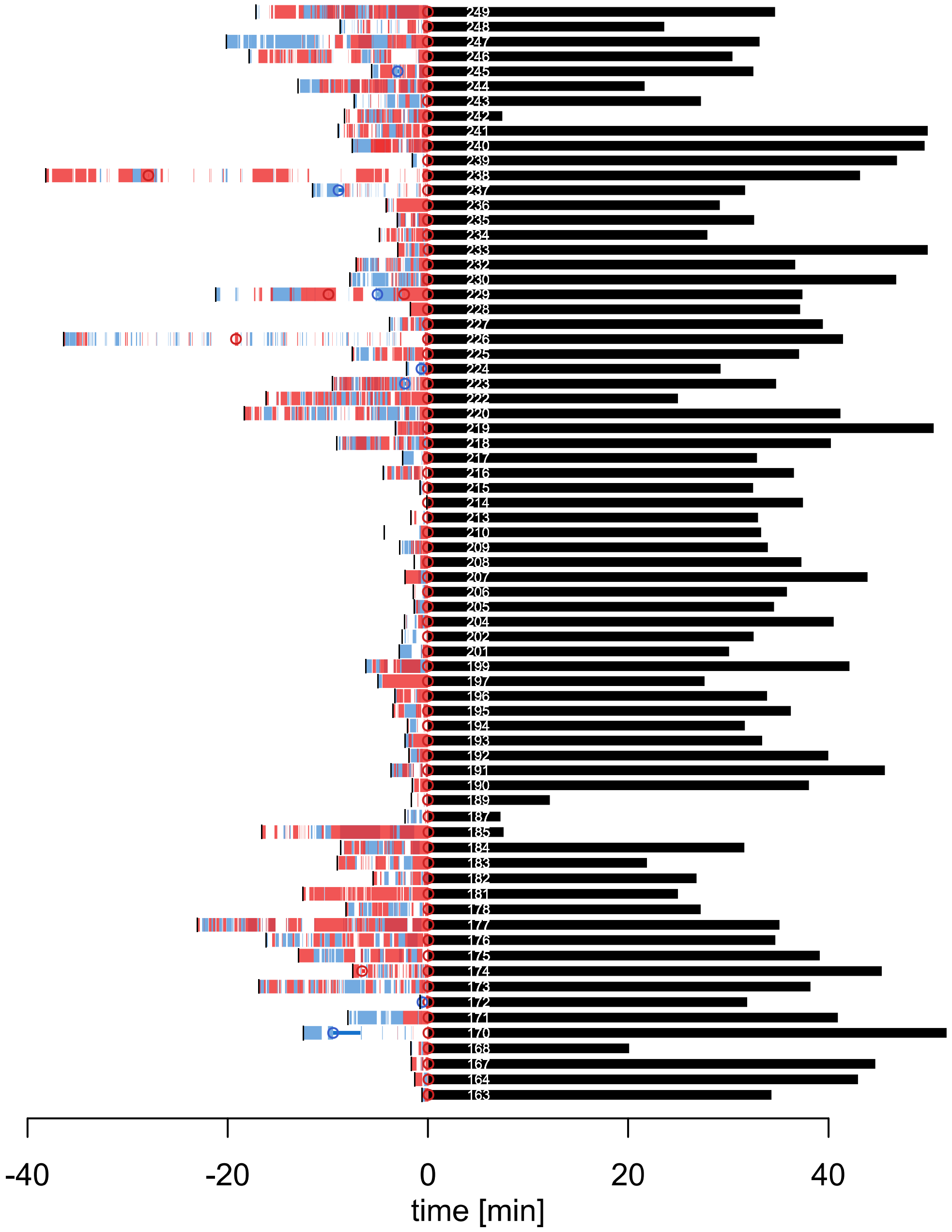


**Figure S2 (continued): Courtship and copulation duration in competition mating experiments.** The black dash indicates courtship start, segments indicate licking periods, open circles mark mounting events and horizontal bars indicate the durations of failed mounting attempts of the copulating male (red) and the non-copulating male (blue). The black bar indicates copulation duration and the number refers to the trial (mating ID), described in Additional datafile 3 and Additional datafile 4.

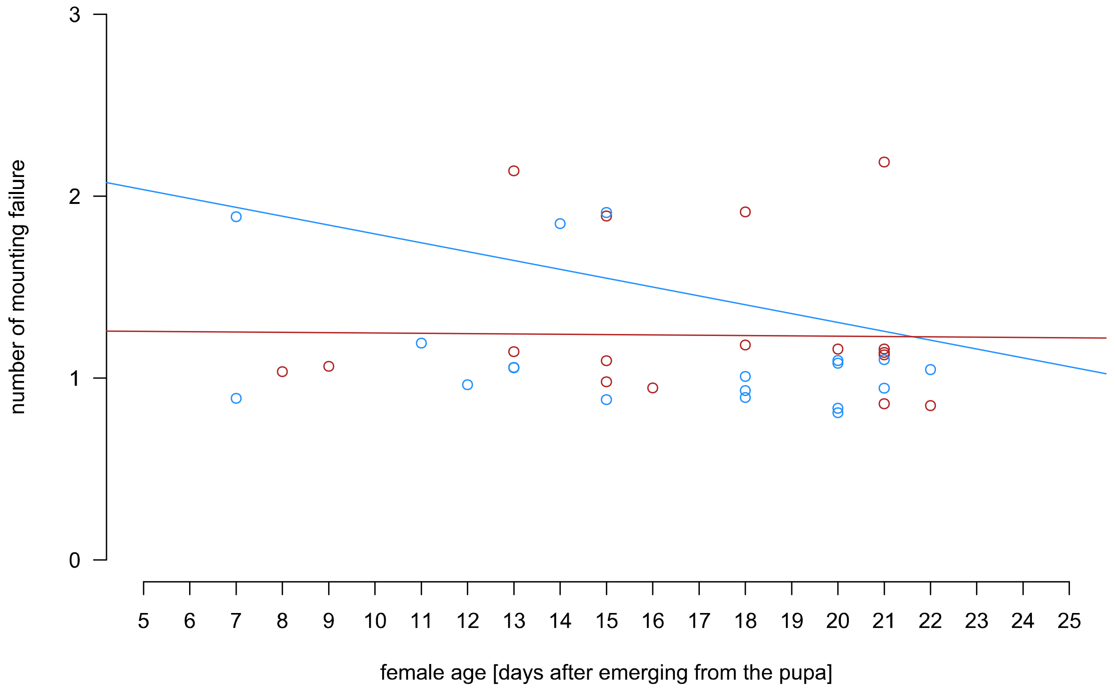


**Figure S3: Failed copulation attempts compared to female age.** Failed copulation attempts of the non-copulating male (blue circles) and the copulating male (red circles) with linear regression lines (non-copulating males: $r^2 = 0.019694$, $y = -0.04872x + 2.2795$, blue line, and copulating males: $r^2 = 0.00029967$, $y = -0.001731x + 1.2643$, red line).

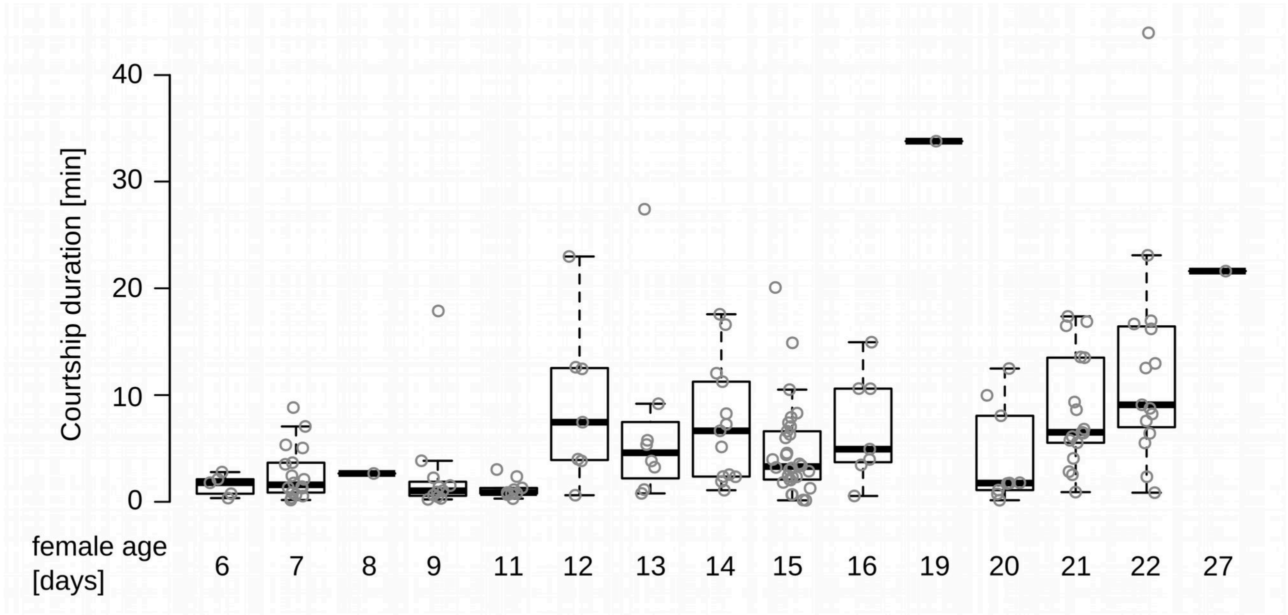


**Figure S4: Courtship duration increases with female age.** The average courtship duration is increased in females that are older than 11 days after hatching from the pupa (Mann-Whitney test, W = 1525.5, *N females younger than 12 days / N females older than 11 days = 52 / 161, P* = $5.811 \times 10^{-12}$). Pearson correlation of courtship duration and female age: 0.41, P= $4.68 \times 10^{-6}$.

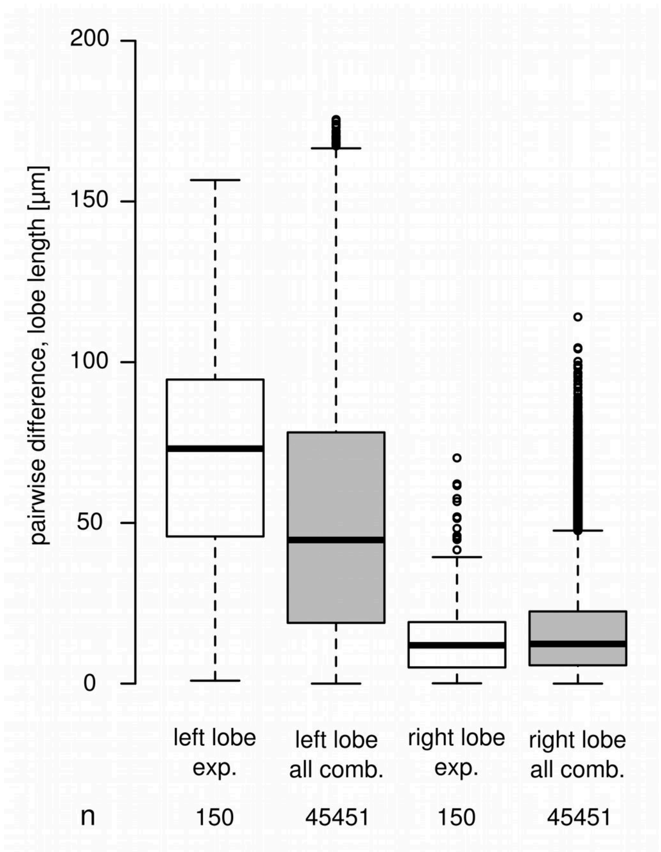


**Figure S5: Pairwise lobe length differences of males used in the competition mating experiments.** The lobe length pairwise length differences of selected pairs in our mating competition experiments (exp., white boxplots) compared to lobe length differences of the same males but randomly compared (random comp., grey boxplots).

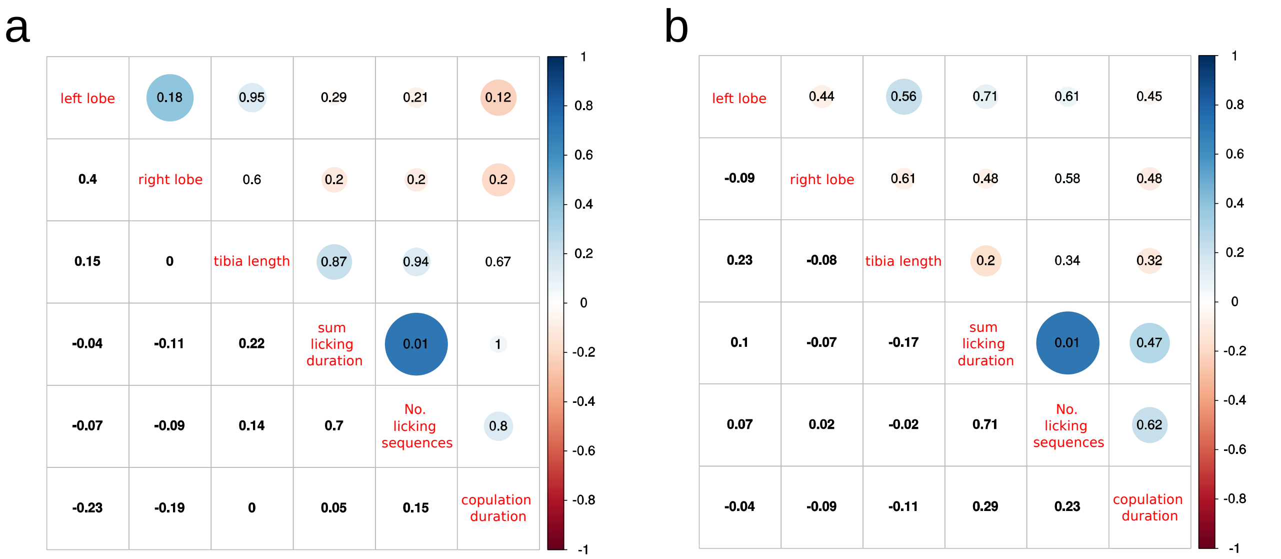


**Figure S6: Pearson correlations of male specific predictor variables. a)** dataset with unmodified males (92 trials), **b)** dataset with lobe-modified males (53 trials). Variable names are listed in the square diagonal, coefficients are in the lower and P-values in the upper half of the square. Circle colors indicate the coefficient according to the scale bars on the right and circle size is proportional to the P-value.

### Table S1: *Drosophila pachea* Resources

| source | stock number | collection locality | collection year |
|---|---|---|---|
| Drosophila Species Stock Center | 15090-1698.01 | USA<br>Arizona<br>Organ Pipe Cactus National Monument | 1997 |
| Drosophila Species Stock Center | 15090-1698.02 | Mexico<br>Sonora<br>Bahia de Kino | 1996 |

### Table S2: Genital lobe lengths in *D. pachea* stocks. †Confidence interval

| stock | left lobe [µm], median (95% CI†) | right lobe [µm] median, (95% CI†) | n |
|---|---|---|---|
| 15090-1698.01 | 206.7 (169.7 - 215.4) | 147.6 (145.4 - 153.1) | 50 |
| 15090-1698.02 | 204.7 (190.5 - 209.2) | 145.4 (138.6 - 146.4) | 50 |
| selection stock | 186.1 (171.8 - 185.0) | 142.2 (138.7 - 144.2) | 99 |

### Table S3: Courtship duration, male licking behavior and copulation duration in trials where both males courted the female

| female | male | n | total courtship duration [min] | | sum licking duration [min] | | No. licking sequences | | copulation duration [min] | |
|---|---|---|---|---|---|---|---|---|---|---|
| | | | median | range | median | range | median | range | median | range |
| 15090-1698.01 | non-copulating sel. lobe length | 47 | 3.23 | 0.30-33.80 | 0.63 | 0.02-7.68 | 4 | 1-22 | --- | --- |
| | copulating sel. lobe length | 47 | | | 1.00 | 0.10-12.53 | 3 | 1-29 | 32.84 | 10.13-49.21 |
| 15090-1698.02 | non-copulating sel. lobe length | 57 | 6.3 | 0.74-23.10 | 1.58 | 0.04-14.24 | 5 | 1-36 | --- | --- |
| | copulating sel. lobe length | 57 | | | 2.80 | 0.14-14.58 | 5 | 1-35 | 27.7 | 2.72-50.21 |
| 15090-1698.01 | non-copulating modif. lobe length | 54 | 7.2 | 0.12-38.18 | 1.28 | 0.02-12.22 | 7 | 1-64 | --- | --- |
| | copulating modif. lobe length | 54 | | | 1.84 | 0.10-14.92 | 6.5 | 1-34 | 33.05 | 3.45-49.23 |

**Table S4: Competition mating trials used for courtship analysis.** †Experiment excluded when: At least one fly escaped, died, or was injured, the movie recording or data storage failed, or genitalia dissection failure (Additional datafile 3).

| female stock | male stock | Lobe surgery | No. Experiments | | | | | |
|---|---|---|---|---|---|---|---|---|
| | | | total | included | excluded | | male courtship | |
| | | | | | no copulation / courtship | other† | one | both |
| 15090-1698.01 | selection stock | No | 98 | 76 | 16 / 4 | 2 | 25 | 51 |
| 15090-1698.02 | selection stock | No | 89 | 76 | 4 / 0 | 9 | 16 | 60 |
| 15090-1698.01 | selection stock | Yes | 62 | 61 | 1 / 0 | 0 | 7 | 54 |

**Table S5: Bradley-Terry Model fit for 92 trials with non lobe-modified males, evaluation of male-specific predictor variables with exclusion of number of licking sequences.** Model: copulation success ~ left lobe length + right lobe length + tibia length + failed copulation attempt + sum licking duration, null deviance: 127.539 on 92 degrees of freedom, residual deviance: 94.643 on 86 degrees of freedom.

| variable | estimate | standard error | z-value | P (>\|z\|) |
|---|---|---|---|---|
| left lobe length | 0.007358 | 0.003509 | 2.097 | **0.035997** |
| right lobe length | -0.016259 | 0.015030 | -1.082 | 0.279366 |
| tibia length | 0.011875 | 0.007138 | 1.664 | 0.096199 |
| failed copulation attempt | -0.347733 | 0.554350 | -0.627 | 0.530476 |
| sum licking duration | 0.769157 | 0.221818 | 3.468 | **0.000525** |

**Table S6: Bradley-Terry Model fit for 92 trials with non lobe-modified males, evaluation of male-specific predictor variables with exclusion of sum licking duration.** Model: copulation success ~ left lobe length + right lobe length + tibia length + failed copulation attempt + number of licking sequences, null deviance: 127.539 on 92 degrees of freedom, residual deviance: 114.96 on 87 degrees of freedom.

| variable | estimate | standard error | z-value | P (>\|z\|) |
|---|---|---|---|---|
| left lobe length | 0.007008 | 0.003175 | 2.207 | **0.0273** |
| right lobe length | -0.005840 | 0.012564 | -0.465 | 0.6420 |
| tibia length | 0.012172 | 0.006625 | 1.837 | 0.0662 |
| failed copulation attempt | -0.071513 | 0.448564 | -0.159 | 0.8733 |
| number of licking sequences | 0.093730 | 0.060826 | 1.541 | 0.1233 |

**Table S7: Bradley-Terry Model fit for 92 trials with non lobe-modified males, evaluation of interactions of male-specific variables with female stock.** Model: copulation success ~ (left lobe length + right lobe length + tibia length + failed copulation attempt + sum licking duration) * female stock, null deviance: 127.539 on 92 degrees of freedom, residual deviance: 87.62 on 82 degrees of freedom.

| variable | estimate | standard error | z-value | P (>\|z\|) |
|---|---|---|---|---|
| left lobe length | 0.0070509 | 0.0053873 | 1.309 | 0.1906 |
| right lobe length | -0.0164323 | 0.0221690 | -0.741 | 0.4586 |
| tibia length | 0.0002137 | 0.0122500 | 0.017 | 0.9861 |
| failed copulation attempt | -0.3510129 | 0.6557440 | -0.535 | 0.5924 |
| sum licking duration | 0.4372115 | 0.2212514 | 1.976 | **0.0481** |
| female stock (1698.02) | NA | NA | NA | NA |
| left lobe length * female stock (1698.02) | 0.0057967 | 0.0086074 | 0.673 | 0.5007 |
| right lobe length * female stock (1698.02) | -0.0144092 | 0.0346259 | -0.416 | 0.6773 |
| tibia length * female stock (1698.02) | 0.0235552 | 0.0166775 | 1.412 | 0.1578 |
| failed copulation attempt * female stock (1698.02) | 0.0355114 | 1.2123092 | 0.029 | 0.9766 |
| sum licking duration * female stock (1698.02) | 1.0801333 | 0.5623220 | 1.921 | 0.0548 |

**Table S8: Bradley-Terry Model fit for 92 trials with non lobe-modified males, evaluation of interactions of male-specific variables with courtship duration.** Sum licking duration and number of licking sequences were excluded from the model because they are redundant with courtship duration. Model: copulation success ~ (left lobe length + right lobe length + tibia length + failed copulation attempt) * courtship duration, null deviance: 127.539 on 92 degrees of freedom, residual deviance: 115.21 on 84 degrees of freedom.

| variable | estimate | standard error | z-value | P (>\|z\|) |
|---|---|---|---|---|
| left lobe length | 0.0046696 | 0.0052433 | 0.891 | 0.373 |
| right lobe length | -0.0038326 | 0.0191598 | -0.200 | 0.841 |
| tibia length | 0.0059164 | 0.0096083 | 0.616 | 0.538 |
| failed copulation attempt | -1.1040192 | 1.0071044 | -1.096 | 0.273 |
| courtship duration | NA | NA | NA | NA |
| left lobe length * courtship duration | 0.0003350 | 0.0006249 | 0.536 | 0.592 |
| right lobe length * courtship duration | -0.0005112 | 0.0021788 | -0.235 | 0.814 |
| tibia length * courtship duration | 0.0008349 | 0.0012409 | 0.673 | 0.501 |
| failed copulation attempt* courtship duration | 0.1451978 | 0.1150052 | 1.263 | 0.207 |

**Table S9: Bradley-Terry Model fit for 92 trials with non lobe-modified males, evaluation of interactions of male-specific variables with female age.** Model: copulation success ~ (left lobe length + right lobe length + tibia length + failed copulation attempt + sum licking duration) * female age, null deviance: 127.539 on 92 degrees of freedom, residual deviance: 87.95 on 82 degrees of freedom.

| variable | estimate | standard error | z-value | P (>\|z\|) |
|---|---|---|---|---|
| left lobe length | 1.520e-02 | 1.274e-02 | 1.193 | 0.2327 |
| right lobe length | -5.233e-02 | 5.401e-02 | -0.969 | 0.3326 |
| tibia length | 1.136e-02 | 2.200e-02 | 0.516 | 0.6057 |
| failed copulation attempt | -4.394e+00 | 2.128e+00 | -2.065 | **0.0389** |
| sum licking duration | 1.190e+00 | 9.719e-01 | 1.225 | 0.2207 |
| female age | NA | NA | NA | NA |
| left lobe length * female age | -4.739e-04 | 7.608e-04 | -0.623 | 0.5334 |
| right lobe length * female age | 2.102e-03 | 3.389e-03 | 0.620 | 0.5350 |
| tibia length * female age | 5.203e-05 | 1.448e-03 | 0.036 | 0.9713 |
| failed copulation attempt * female age | 2.773e-01 | 1.322e-01 | 2.098 | **0.0359** |
| sum licking duration * female age | -1.624e-02 | 5.514e-02 | -0.294 | 0.7684 |

**Table S10: Bradley–Terry (BT) model examining the effects of left lobe length, sum licking duration and failed mounting attempts on copulation success,** in 92 trials with unmodified males. The main effect for “female age” cannot be estimated in the BT model because it is specific to each trial and not to each male. Bolded estimates are significant. Model: copulation success ~ sum licking duration + left lobe length + failed mounting attempts * female age, null deviance: 127.539 on 92 degrees of freedom, residual deviance: 92.813 on 88 degrees of freedom. Influential trials 29 and 141 were detected to affect model estimates.

| variable | estimate | standard error | z-value | P (>\|z\|) |
|---|---|---|---|---|
| sum licking duration | 0.867081 | 0.244225 | 3.550 | **0.000385** |
| left lobe length | 0.006994 | 0.003450 | 2.027 | **0.042637** |
| failed copulation attempt | -3.7406217 | 1.783746 | -2.097 | **0.035989** |
| female age | NA | NA | NA | NA |
| failed mounting attempts x female age | 0.245507 | 0.117685 | 2.086 | **0.036966** |

**Table S11: Bradley–Terry (BT) model examining the effects of left lobe length, right lobe length, number of licking sequences, tibia length and failed mounting attempts on copulation success,** in 53 trials with lobe-modified males.Model: copulation success ~ left lobe length + right lobe length + tibia length + failed copulation attempt + number of licking sequences, null deviance: 73.474 on 53 degrees of freedom, residual deviance: 69.250 on 48 degrees of freedom.

| variable | estimate | standard error | z-value | P (>\|z\|) |
|---|---|---|---|---|
| left lobe length | 0.0088769 | 0.0048547 | 1.829 | 0.0675 |
| right lobe length | 0.0098511 | 0.0229071 | 0.430 | 0.6672 |
| tibia length | -0.0007568 | 0.0079174 | -0.096 | 0.9239 |
| failed copulation attempt | 0.0611166 | 0.8192932 | 0.075 | 0.9405 |
| number of licking sequences | -0.0340544 | 0.0412683 | -0.825 | 0.4093 |

**Table S12: Bradley–Terry (BT) model examining the effects of left lobe length, right lobe length, sum licking duration, tibia length and failed mounting attempts on copulation success,** in 53 trials with lobe-modified males. Model: copulation success ~ left lobe length + right lobe length + tibia length + failed copulation attempt + sum licking duration, null deviance: 73.474 on 53 degrees of freedom, residual deviance: 67.020 on 48 degrees of freedom.

| variable | estimate | standard error | z-value | P (>\|z\|) |
|---|---|---|---|---|
| left lobe length | 0.0067083 | 0.0049313 | 1.360 | 0.174 |
| right lobe length | 0.0064819 | 0.0234845 | 0.276 | 0.783 |
| tibia length | 0.0009478 | 0.0079822 | 0.119 | 0.905 |
| failed copulation attempt | -0.3486010 | 0.9209974 | -0.379 | 0.705 |
| sum licking duration | 0.2443697 | 0.1641949 | 1.488 | 0.137 |
